# RAD51AP1 mediates RAD51 activity through nucleosome interaction

**DOI:** 10.1101/2020.12.17.421636

**Authors:** Elena Pires, Neelam Sharma, Platon Selemenakis, Bo Wu, Yuxin Huang, Dauren S. Alimbetov, Weixing Zhao, Claudia Wiese

## Abstract

RAD51 Associated Protein 1 (RAD51AP1) is a key protein in the homologous recombination DNA repair pathway (HR). Loss of RAD51AP1 leads to defective HR, genome instability and telomere erosion. RAD51AP1 physically interacts with the RAD51 recombinase and promotes RAD51-mediated capture of the donor DNA, synaptic complex assembly and displacement-loop formation when tested with synthetic, nucleosome-free DNA substrates *in vitro*. In cells, however, DNA is packaged into chromatin, posing an additional barrier to the complexities of the HR reaction. How RAD51AP1 functions as an HR activator in the context of chromatin has remained unclear.

In this study, we show that RAD51AP1 binds to Nucleosome Core Particles (NCPs). We identified a C-terminal region in RAD51AP1 and its previously mapped DNA binding domain as critical for mediating the association between RAD51AP1 and both the NCP and the histone octamer. We show that RAD51AP1 is capable of promoting duplex DNA capture and initiating joint-molecule formation with the NCP and chromatinized template DNA, respectively. Together, our results suggest that RAD51AP1 *directly* assists the RAD51-mediated search of donor DNA in chromatin. We present a model, in which RAD51AP1 anchors the DNA template through affinity for its nucleosomes to the RAD51-ssDNA nucleoprotein filament.

## Introduction

Homologous recombination DNA repair (HR) is a template-dependent DNA damage repair pathway critical for genome stability and cancer avoidance. HR also is essential for the smooth progression of DNA replication (1). Central to the HR reaction is the RAD51-ssDNA nucleoprotein filament, also called the presynaptic filament, which captures the DNA donor, engages in synapsis and generates a displacement-loop (D-loop) upon location of the homologous DNA target sequence.

RAD51AP1 is an intrinsically unfolded protein (2), and is likely to undergo induced folding upon binding to specific partners which will then make it well-ordered (3). RAD51AP1 binds to the RAD51 recombinase and stimulates RAD51-mediated joint-molecule formation, as we and others have shown (4–10). In RAD51AP1, two distinct DNA binding domains have been identified (4,6). Both domains are required for the full activity of RAD51AP1 in protecting cells from DNA damaging agents and in joint-molecule formation assays with synthetic DNA substrates *in vitro* (6).

Previous efforts to elucidate the biochemical properties of RAD51AP1 relied on nucleosome-free DNA substrates (4–9). In eukaryotic cells, however, DNA is packaged into chromatin with nucleosomes forming the minimum basic unit. How RAD51AP1 would promote RAD51 activity in the context of reconstituted nucleosome core particles (NCPs) and nucleosome-containing donor DNA was unclear.

In this study, we show that RAD51AP1 physically associates with the NCP. Binding to the NCP retains the RAD51AP1-RAD51 interaction. In the context of the NCP, RAD51AP1 stimulates the capture of the DNA template. RAD51AP1 also stimulates RAD51-mediated strand invasion into chromatinized DNA. Collectively, our results suggest that RAD51AP1 is an important accessory HR factor in the chromatin environment and functions as a bridging molecule between the RAD51-ssDNA nucleoprotein filament and the incoming homologous chromatin donor. We also show that a previously identified DNA binding domain in RAD51AP1 is critical for establishing the contact with the NCP and strand invasion.

## Results

### RAD51AP1 interacts with the mono-Nucleosome Core Particle (NCP)

Intrigued by our earlier findings that showed affinity of purified human RAD51AP1 to chromatinized DNA in the immobilized template assay (11), we decided to test complex formation between RAD51AP1 and the NCP. The NCP is the minimum basic unit of chromatin in which ∼2 super-helical turns of 147 bp double-stranded (ds)DNA are wrapped around one histone octamer with no free DNA ends remaining (12,13). We reconstituted NCPs by salt gradient deposition using the 601 Widom 147 bp dsDNA fragment and human histone octamers, as previously described (12,14), and used electrophoretic mobility shift assays (EMSAs) to assess the RAD51AP1-NCP association (for schematic of the assay see Fig. 1*A*). Following incubation of full-length human His_6_-tagged RAD51AP1 (Fig. S1*A*) with the NCP, we observed the identical RAD51AP1-mediated mobility shifts by ethidium bromide first, and then by Imperial protein stain (Fig. 1*B*, *lanes 4* and *5*). Of note, Imperial protein stain does not stain DNA (Fig. S1*B*, *lanes 1* and *2*). As determined by Western blot analysis, RAD51AP1 enters the native gel in the presence but not in the absence of the NCP (Fig. 1*B*, compare *lanes 6* and *7* to *lane 8*), further supporting complex formation. We also expressed and purified His_6_-/FLAG-tagged human RAD51AP1 (Fig. S1*C*) and detected NCP binding activity for this protein as well (Fig. 1*C*, *lanes 3* and *4*, and *D*, *lanes 2* and *3*). Moreover, RAD51AP1 protein and both histone H2A and histone H3 were detected within the mobility shifted bands (Fig. 1*C*, *lanes 5*-*10*, and *D*, *lanes 4*-*9*), suggestive of a tight RAD51AP1/NCP complex inclusive of octamer components. We also monitored NCP binding of purified human FLAG-tagged RAD54 (Fig. S1*D*), a member of the SWI2/SNF2 group of ATP-dependent chromatin-remodeling factors previously shown to efficiently bind to a core nucleosome with no DNA overhangs (15,16). We found similar activities for both RAD51AP1 and RAD54 in NCP binding (Fig. 1*E*), further supporting the validity of our conducted experiments and obtained results. Purified human RAD51 (Fig. S1*E*), however, showed no affinity for the NCP, as determined by EMSA (Fig. 1*F*).

**Figure 1.**
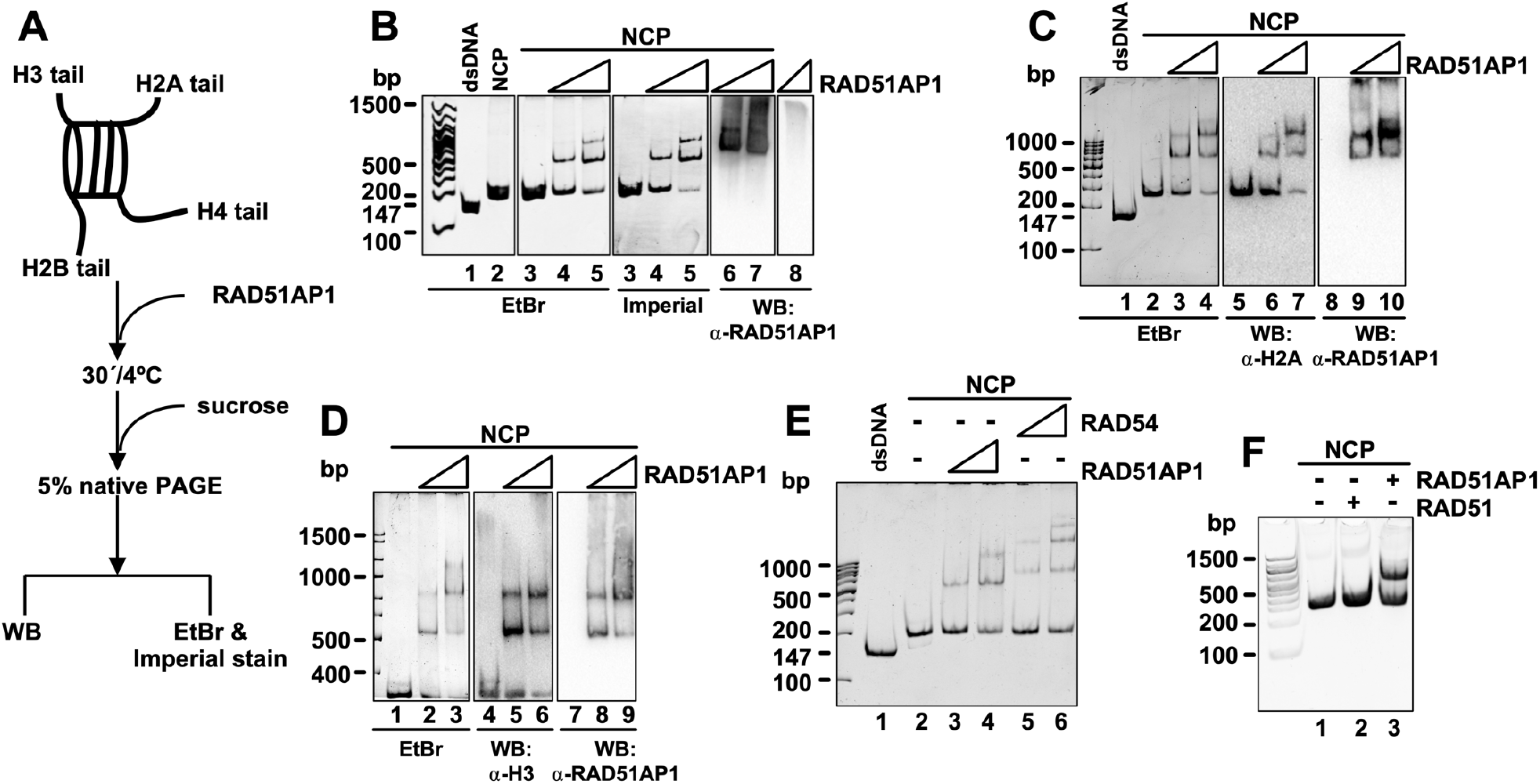
RAD51AP1 interacts with the Nucleosome Core Particle (NCP). ***A***, Schematic of the NCP and the EMSA protocol. ***B***, EMSA of RAD51AP1-His_6_ (0.1 and 0.2 μM) with the NCP (0.2 μM) assessed by ethidium bromide (lanes 3-5) first and then by Imperial protein stain (lanes 3-5). The Western blots show the presence of RAD51AP1 entering the gel only when bound to the NCP (lanes 6-7), not alone (lane 8). ***C***, EMSA of His_6_/FLAG-tagged RAD51AP1 (0.4 and 0.8 μM) with the NCP (0.4 μM) visualized by ethidium bromide (lanes 1-4). A second gel was loaded in duplicate, transferred and probed to histone H2A or RAD51AP1. ***D***, EMSA of His_6_/FLAG-tagged RAD51AP1 (0.4 and 0.8 μM) with the NCP (0.4 μM) visualized by ethidium bromide (lanes 1-3). A second gel was loaded in duplicate, transferred and probed to histone H3 or RAD51AP1. ***E***, EMSA of His6/FLAG-tagged RAD51AP1 (0.1 and 0.2 μM) and of FLAG-tagged RAD54 (0.1 and 0.2 μM) with the NCP (0.2 μM) assessed by ethidium bromide (lanes 3-4 and 5-6, respectively). ***F***, RAD51 does not bind to the NCP. EMSA of RAD51 and RAD51AP1 (0.4 and 0.2 μM, respectively) with the NCP (0.2 μM).

During DNA repair, nucleosomes are dynamically assembled and disassembled, guided by histone chaperones and ATP-dependent chromatin remodeling factors (17), and several DNA repair proteins with histone chaperone activity have been identified (18–21). To test for the possibility that RAD51AP1 may function as a histone chaperone, we used a nucleosome reconstitution assay and purified human NAP1 (Fig. S1*F*), a protein with well-established histone chaperone activity (22–24), as a positive control. Our results show that, unlike NAP1, RAD51AP1 does not possess histone chaperone activity (Fig. S1*G*, *lanes 4-6*, and *H*, *lanes 4-8*).

### The C-terminal domain of RAD51AP1 is sufficient for NCP interaction

After confirming the novel association between RAD51AP1 and the NCP, we identified the domain in RAD51AP1 responsible for complex formation. We purified three previously described (6–8), non-overlapping MBP-/His_6_-tagged fragments of the RAD51AP1 protein (F1, F2 and F3) (see Fig. 2*A* for schematic; Fig. S2*A*) and also full-length MBP-/His_6_-RAD51AP1 (Fig. S2*B*). RAD51AP1-F1 and -F3 each contain one mapped DNA binding domain critical for protein function, as previously reported and identified using nucleosome-free DNA (6). We found that RAD51AP1-F3 is the only fragment capable of binding to the NCP (Fig. 2*B*, *lane 7*), whereas RAD51AP1-F1 and RAD51AP1-F2 are devoid of this activity (Fig. 2*B*, *lane 5*, and Fig. S2*C*, *lanes 5* and *6*, and *D, lanes 5* and *6*). Complex formation between RAD51AP1-F3 and the NCP was confirmed by gel filtration (Fig. S2*E* and *F*). To determine the apparent binding affinities (*K*_D(app)_) of full-length RAD51AP1 to the 147 bp dsDNA fragment and the NCP, we performed EMSAs and quantified the free dsDNA or free NCP bands, respectively, after ethidium bromide staining (Fig. S2*H* and *I*). After correcting for the diminished ability of ethidium bromide to intercalate into NCP-bound DNA (Fig. S2*J* and *K*), we found the respective composite affinities of full-length RAD51AP1 to be exceptionally tight to dsDNA (*K*_D(app)_ = 12 nM) and the NCP (*K*_D(app)_ = 21 nM).

**Figure 2.**
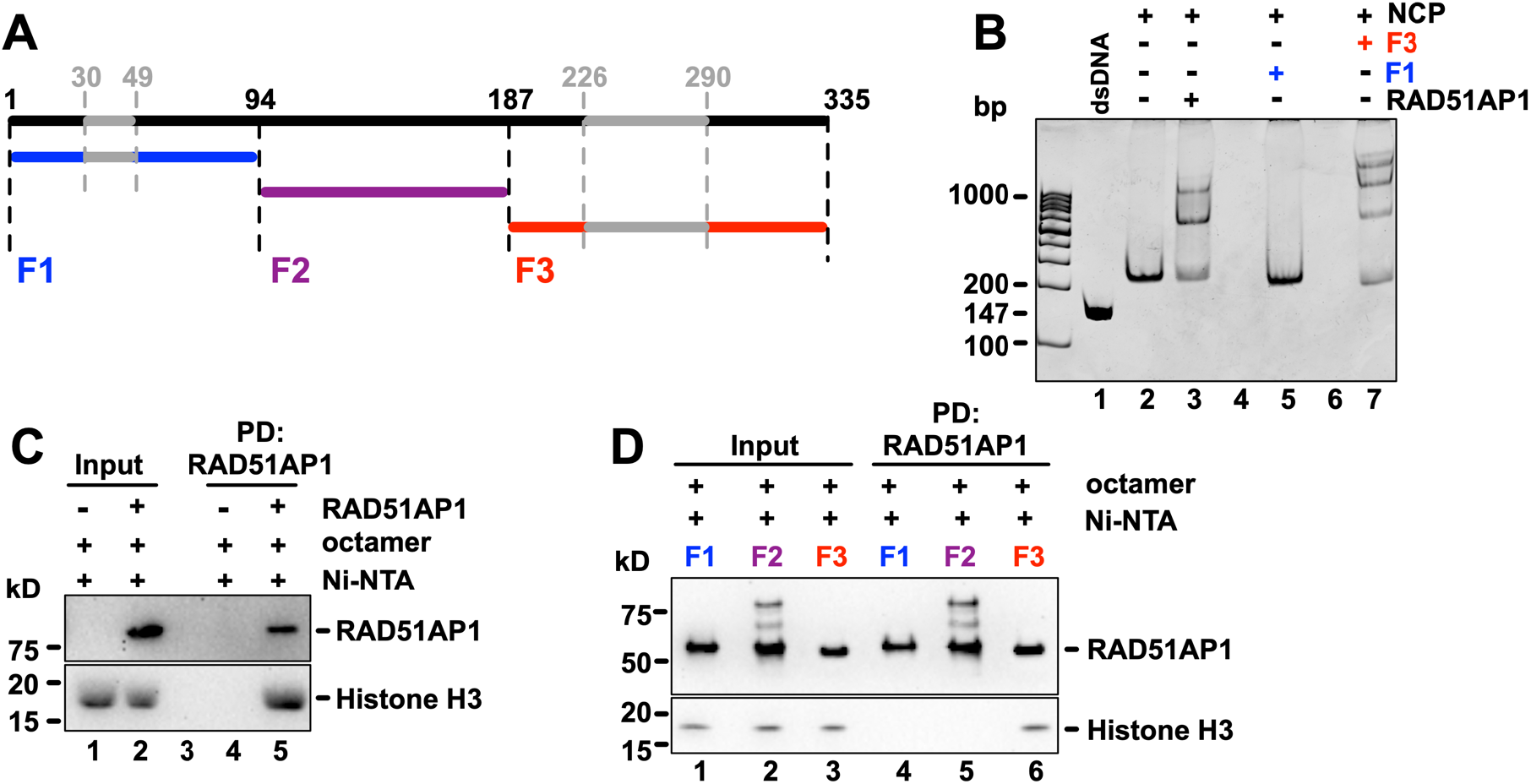
The association between the NCP and RAD51AP1 occurs through the RAD51AP1-F3 domain. ***A***, Schematic of full-length RAD51AP1 (isoform 2; in black) and RAD51AP1 fragments (F1 (blue), F2 (purple) and F3 (red)), as described earlier (7), including the two mapped DNA binding domains (grey), as previously shown (4,6). ***B***, EMSA of MBP-/His_6_-tagged full-length RAD51AP1, -F1 and -F3 (0.2 μM each) with the NCP (0.2 μM) assessed by ethidium bromide. ***C***, Western blots obtained after Ni-NTA pulldown of MBP-RAD51AP1-His_6_ to show that RAD51AP1 *directly* interacts with histone H3 (lane 5). ***D***, Western blots obtained after Ni-NTA pulldown of the three MBP-/His_6_-tagged RAD51AP1 fragments to show that the interaction between RAD51AP1 and histone H3 occurs through the F3 domain (lane 6).

We assessed the possibility of a direct interaction between RAD51AP1 and the histone octamer (Fig. S2*G*) in affinity pulldown assays. We obtained evidence of a direct interaction between full-length RAD51AP1 and RAD51AP1-F3 with histone H3 (Fig. 2*C*, *lane 5* and *D*, *lane 6*, respectively). In contrast, neither RAD51AP1-F1 nor RAD51AP1-F2 interact with histone H3 (Fig. 2*D*, *lanes 4* and *5*).

Further division of the RAD51AP1-F3 fragment and its DNA binding domain into an N-terminal 88 amino acid (aa) residues encompassing fragment (*i.e.*, N88) and a C-terminal 60 aa residues containing fragment (*i.e.*, C60) (see Fig. 3*A* for schematic; Fig. S3*A* and *B*) impaired DNA binding, as expected ((6); Fig. S3*C*, *lanes 4*-*7*). Moreover, neither C60 nor N88 were able to form a complex with the NCP (Fig. 3*B*, *lanes 5* and *6*, and S3*D*, *lanes 6*-*8*). Similarly, RAD51AP1-F3 in which several consecutive lysine residues and a tryptophan, previously and here deemed critical for binding to nucleosome-free DNA ((6); Fig. S3*E-H* and S3*I*, *lanes 5-7* and data not shown), were changed to alanine, lost or showed greatly reduced ability to bind to the NCP (Fig. 3*C*, *lanes 4* and *5*, and *S3J, lanes 4-9*). Collectively, these results show that the previously identified RAD51AP1 DNA binding domain that is located in the F3 fragment (6) mediates complex formation between RAD51AP1 and the NCP, and that contact is made by RAD51AP1 with histone H3 and, likely, nucleosomal DNA.

**Figure 3.**
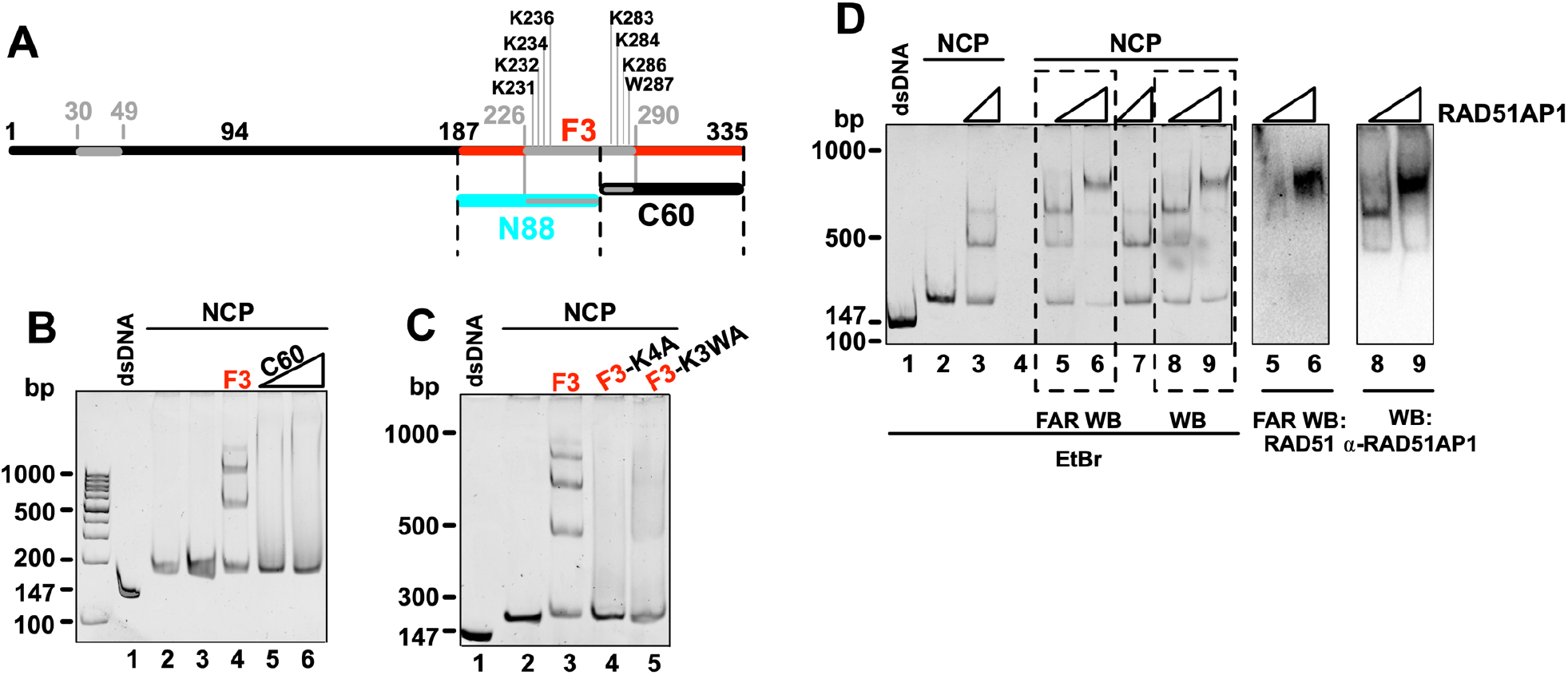
The association between the NCP and RAD51AP1-F3 involves the DNA binding domain. ***A***, Schematic of full-length RAD51AP1 (isoform 2; black), including the RAD51AP1-F3 (red), -N88 (turquoise) and -C60 (black) fragments with the mapped bi-partite DNA binding domain (grey) and its critical residues (*K231, K232, K234, K236, K283, K284, K286 and W287), as previously identified (4,6). ***B***, EMSA of F3 (0.5 μM) and C60 (0.5 and 1 μM) with the NCP (0.4 μM). ***C***, EMSA of F3 (0.5 μM) and DNA bindingdefective mutants of F3 (F3-K4A and F3-K3WA; 1.0 μM each) with the NCP (0.4 μM). K4A: compound point mutant of K231A, K232A, K234A and K236A; K3WA: compound point mutant of K283A, K284A, K286A and W287A. *Residue numbering based on RAD51AP1 isoform 2. ***D***, Western blots and FAR Western analysis after EMSA of full-length His_6_-/FLAG-tagged RAD51AP1 (0.25, 0.5 and 1.0 μM) with the NCP (0.4 μM) to visualize the presence of RAD51AP1 in shifted bands (lanes 8 and 9) and bound RAD51 to the RAD51AP1-NCP complex (lane 6).

### The RAD51AP1-NCP complex retains the capability to bind to RAD51

Since our results confirmed that complex formation between RAD51AP1 and the NCP stems from the F3 region and since the F3 region contains the RAD51 interaction domain (4,5,25), we tested if full-length RAD51AP1 in the RAD51AP1/NCP complex was capable of retaining the interaction with the RAD51 recombinase. To this end, we conducted a FAR Western experiment in which we first separated the RAD51AP1/NCP complex by EMSA, transferred proteins to a PVDF membrane, and then incubated the membrane with purified human RAD51 protein before detection of membrane-bound RAD51 by Western blot analysis. As described previously and shown in Fig. 1, RAD51AP1 is present in shifted bands (Fig. 3*D*, *lanes 8 and 9*). In addition, RAD51 is bound and detected at the super-shifted RAD51AP1/NCP band (Fig. 3*D*, *lane 6*). As shown above, RAD51 by itself is unable to associate with the NCP (Fig. 1*F*). Collectively, these results suggest that the RAD51AP1 protein, while in complex with the NCP, retains the ability to interact with the RAD51 recombinase.

### RAD51AP1 promotes capture of the NCP in the duplex capture assay

Earlier reports on RAD51AP1 showed its ability to stimulate duplex DNA capture in the context of nucleosome-free, naked dsDNA (8,9,26,27). The results presented in this study show that RAD51AP1 interacts with the NCP and that the RAD51AP1/NCP complex retains the ability to interact with RAD51. These findings prompted us to test if RAD51AP1 would stimulate RAD51 activity in the duplex capture assay with an NCP. We tested the ability of RAD51AP1 to capture nucleosome-containing homologous template DNA via the NCP (see Fig. 4*A* for schematic of the assay). In accordance with our earlier findings (8,9,26,27), addition of RAD51AP1 to the reaction stimulated the capture of nucleosome-free dsDNA (Fig. 4*B*, *lanes 3* and *4*). Interestingly, addition of RAD51AP1 also stimulated capture of the NCP (Fig. 4*B*, *lanes 7* and *8*). Quantitative analyses from three independent experiments shows that RAD51AP1 prefers the NCP over nucleosome-free DNA in the duplex capture assay (Fig. 4*C*, *p*<0.05 and *p*<0.01; two-way ANOVA).

**Figure 4.**
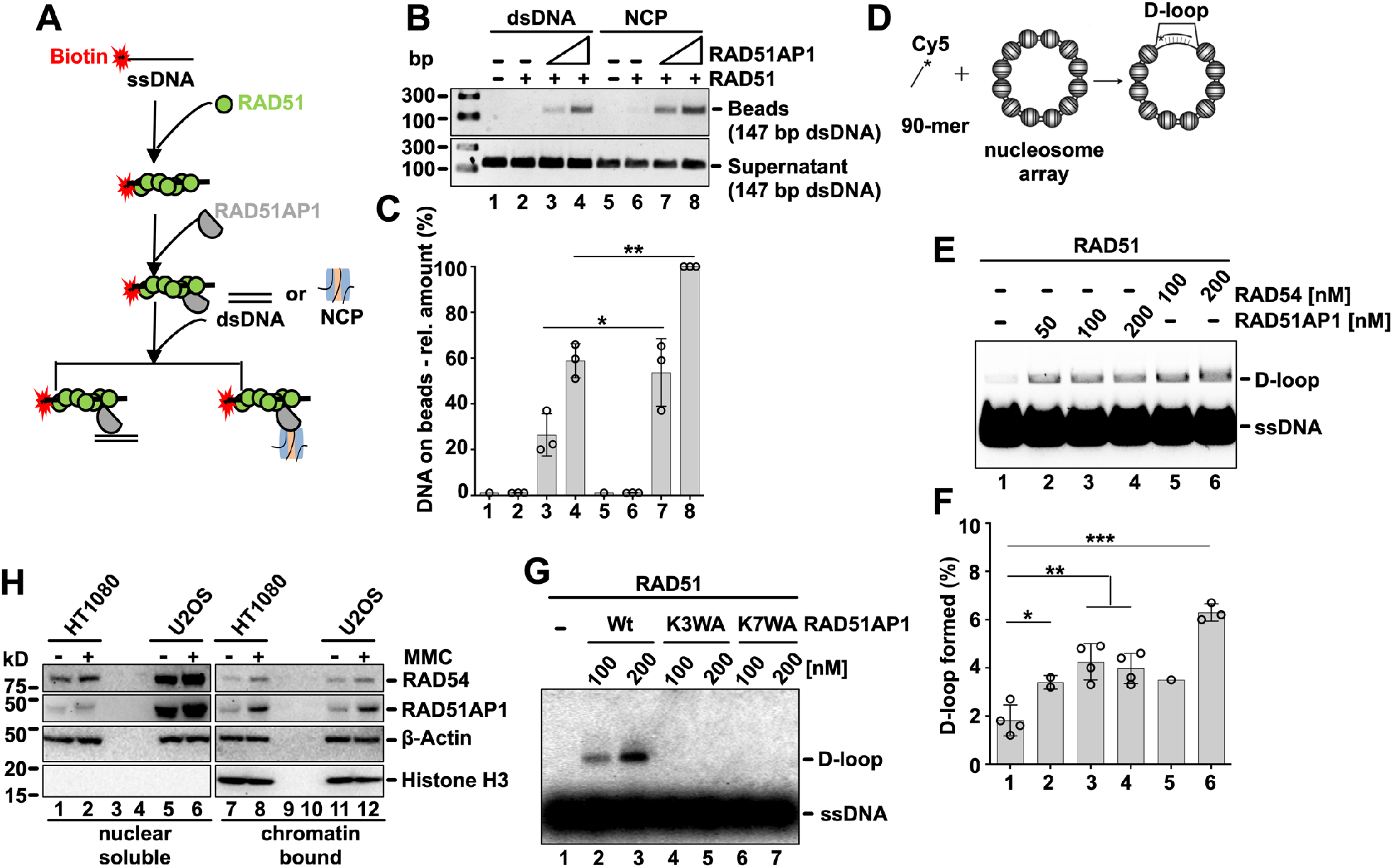
RAD51 binds to the RAD51AP1/NCP complex, RAD51AP1 stimulates duplex capture with the NCP and is recruited to the chromatin fraction in human cells after DNA damage. ***A***, Schematic of the duplex capture assay with His_6_/FLAG-tagged RAD51AP1 and either nucleosome-free DNA (*i.e.*, 147 bp dsDNA) or the NCP. ***B***, Qualitative analysis of captured DNA (beads) and DNA in supernatant by agarose gel electrophoresis. ***C***, Quantitative analysis: Symbols are the results from independent experiments. Bars are the means from three independent experiments ± 1 SD; *, *P* < 0.05; **, *P* < 0.01; two-way ANOVA. ***D***, Schematic of the D-loop reaction with chromatinized pBluescript II SK (-) plasmid DNA. ***E***, Addition of RAD51AP1 (50, 100 and 200 nM; lanes 2-4) or RAD54 (100 and 200 nM; lanes 5-6) promotes the RAD51-mediated Dloop reaction on chromatinized DNA. ***F***, Quantification of the results. Symbols are the results from independent experiments. Bars are the means from 2-4 independent experiments ± 1SD. *, *P* < 0.05; **, *P* < 0.01; ***, *P* < 0.001; multiple *t*-test analysis. ***G***, Agarose gel to show that wild type RAD51AP1 (100 and 200 nM; lanes 2-3) promotes the RAD51-mediated D-loop reaction on chromatinized DNA, but RAD51AP1-K3WA (100 and 200 nM; lanes 4-5) and RAD51AP1-K7WA (100 and 200 nM; lanes 6-7) are unable to do so. ***H***, Western blots of fractionated extracts of the nuclei from HT1080 and U2OS cells without and after exposure to mitomycin C (MMC). The signals for β-Actin and histone H3 serve as loading and fractionation control, respectively. RAD54 is shown for comparison purposes.

### RAD51AP1 promotes strand invasion of a chromatinized DNA template

Following duplex DNA capture, the RAD51 filament engages in synapsis and generates a displacement-loop (D-loop) by invasion of the homologous target sequence. RAD51-mediated strand invasion and the impact of accessory proteins on this activity can be measured *in vitro* by the oligonucleotide-based D-loop assay (28). In this assay, RAD51AP1 was previously shown to be a strong stimulator of heteroduplex DNA formation with nucleosome-free plasmid DNA (4–6,9). Given RAD51AP1’s affinity for the NCP, we were curious to test its stimulation of RAD51-mediated strand invasion on chromatinized template DNA (for schematic of the assay see Fig. 4*D*). We prepared the chromatinized pBluescript II SK(-) donor plasmid (Fig. S4*A*), and, in parallel to RAD51AP1, also assessed RAD54, a known stimulator of heteroduplex DNA joint formation with chromatinized target DNA (15,29–31). Addition of 50-200 nM RAD51AP1 to this reaction stimulated D-loop formation ∼2-fold (Fig. 4*E* and *F*, *lanes 2-4*; *p*<0.05 and *p*<0.01; multiple *t*-test analysis). Addition of 100 and 200 nM RAD54 stimulated D-loop formation ∼2- and 3.5-fold, respectively (Fig. 4*E* and *F*, *lanes 5-6*). We surmise that RAD51AP1 is able to stimulate RAD51-mediated joint-molecule formation with a chromatinized DNA template, although less efficiently than the RAD54 DNA motor protein.

Next, we tested if mutants of the F3 region in RAD51AP1 that abrogate the interaction with the NCP (Fig. 3*C*), would interfere with stimulating RAD51-mediated strand invasion into the chromatinized donor DNA. To this end we constructed two compound mutants in full-length RAD51AP1 (RAD51AP1-K3WA and RAD51AP1-K7WA; for schematic see Fig. S4*B*), purified these mutants (Fig. S4*C*) and tested their activity in the D-loop assay with chromatinized DNA. We found that both RAD51AP1-K3WA and -K7WA are completely defective in stimulating strand invasion into chromatin (Fig. 4*G*, *lanes 4-7*, and S4*E*, *lanes 5-7*, and *F, lanes 5-7*), although both mutants retained the ability to stimulate strand invasion into naked, nucleosome-free plasmid DNA (Fig. S4*H*, *lanes 4-7*). Together, these results provide strong evidence that the F3 domain in RAD51AP1 is critical for guiding homology search and joint-molecule formation within the context of chromatin.

### RAD51AP1 is recruited to the chromatin fraction in human cells after induced DNA damage

To assess the movement of nuclear soluble RAD51AP1 to the chromatin fraction in human cells, we fractionated nuclear extracts of HT1080 and U2OS cells that were grown under normal conditions or exposed to exogenous DNA damage by treatment with mitomycin C (MMC), an inter-strand crosslinking agent that challenges HR DNA repair. Supporting our earlier findings obtained in HeLa cells (11), little RAD51AP1 is localized to the chromatin in HT1080 or U2OS cells cultured under normal growth conditions (Fig. 4*H*, *lanes 7* and *11*). However, upon exposure to MMC, increased amounts of RAD51AP1 protein are detected at the chromatin fraction in both HT1080 and U2OS cells (Fig. 4*H*, *lanes 8* and *12*). These results show that some activity of RAD51AP1 function is associated with the chromatin fraction in both HT1080 and U2OS cells.

## Discussion

HR involves the exchange of genetic information between homologous DNA molecules and is essential for preserving genome stability (1,32,33). The central steps of the HR reaction are the search for and the identification of a homologous DNA donor sequence, and the formation of a joint molecule catalyzed by the RAD51 recombinase. Based on studies with nucleosome-free synthetic DNA substrates (4–9), the RAD51 recombinase is assisted in this process by the accessory RAD51AP1 protein, for which a role in bridging or anchoring of the two molecules undergoing exchange has been discussed (4,5,9,34). Yet, how exactly these steps are orchestrated in the chromatin environment in cells remains poorly understood.

Intrigued by our previous findings that showed binding of RAD51AP1 to a chromatinized DNA substrate *in vitro* (11), we were eager to test the affinity of RAD51AP1 to an NCP. To avoid any complications from direct binding of RAD51AP1 to overhanging free dsDNA, the NCPs that we used here were free of any linker DNA extensions. We show that RAD51AP1 avidly binds to the NCP and that complex formation is mediated *via* a C-terminal domain in RAD51AP1, a domain with previously identified DNA binding activity (4,6). In contrast, RAD51AP1-F1 which contains the second and N-terminally located DNA binding domain has no ability in complex formation with the NCP. We also discovered a direct physical interaction between RAD51AP1-F3 and histone H3. Accordingly, we suggest that the RAD51AP1/NCP interface could be composed of RAD51AP1 residues in contact with the nucleosomal DNA and with histone H3, although other regions of contact cannot be excluded at this point. In our preferred interpretation of the results obtained, we suggest that the C-terminally located DNA binding domain (in RAD51AP1-F3) associates with the nucleosomes of the homologous dsDNA target, while both the N-terminally located DNA binding domain (in RAD51AP1-F1) and the peptide motif facilitating RAD51 interaction (25) associate with the presynaptic filament. As such, both DNA molecules may be tethered together to prepare for the exchange (Fig. 5). This model would be similar to the mechanisms of dsDNA capture proposed for Hop2-Mnd1 in meiotic recombination in yeast (35).

**Figure 5.**
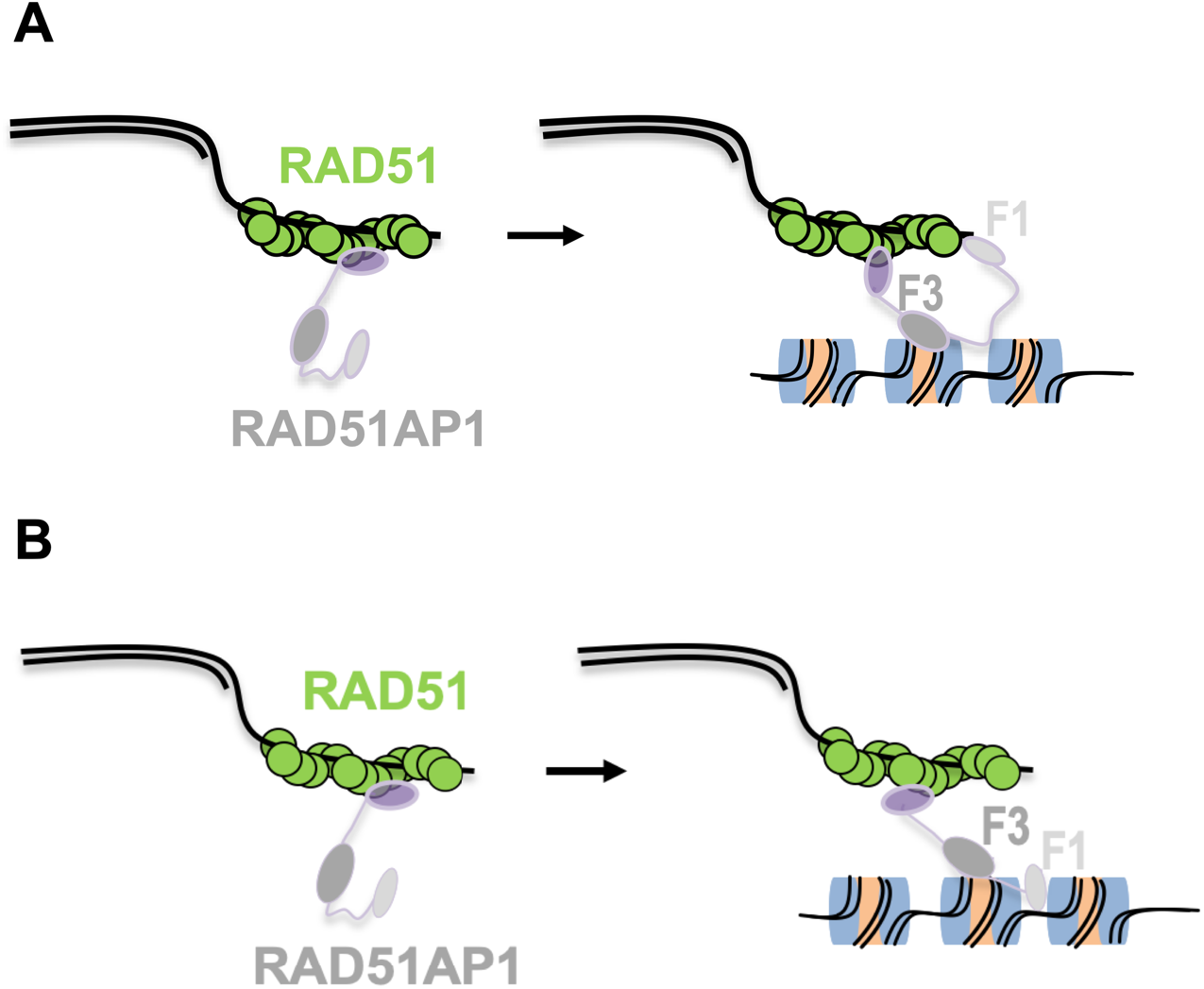
Model depicting the role of the RAD51AP1-F1 and -F3 domains in ternary complex formation. ***A***, To initiate ternary complex formation, F3 (dark grey) associates with a nucleosome on the incoming duplex DNA template, while F1 (light grey) binds to the ssDNA of the RAD51-ssDNA nucleoprotein filament. The very C-terminus of RAD51AP1 (purple) engages with RAD51, as previously shown (25). ***B***, It is also possible that F1 binds to regions of nucleosome-free DNA within the dsDNA target.

We detected at least two major shifted species when RAD51AP1 was incubated with the NCP. These observations suggest multiple contacts between RAD51AP1 and the nucleosome. It is possible that one species is mediated predominantly through contact the with the nucleosomal DNA, while the second species associates predominantly through contacting histones. Further assessing the RAD51AP1:NCP stoichiometry in the future will be important to determine if two RAD51AP1 molecules can occupy one single nucleosome.

A select few proteins function synergistically with RAD51AP1 in biochemical HR assays based on synthetic nucleosome-free DNA substrates and also in cells. For example, RAD51AP1 interacts with UAF1 and the RAD51AP1-UAF1 complex works in conjunction with the RAD51 filament in duplex capture and synaptic complex assembly (26). As in RAD51AP1, a DNA binding domain in UAF1 was identified that is critically important for HR-mediated chromosome damage repair (27,36). Hence, it will be important to test if UAF1 shows affinity for nucleosome-containing DNA, and if UAF1 can synergize with RAD51AP1 in the homologous pairing reaction with chromatin substrates. Similarly, functional synergism also was determined between RAD51AP1 and the tumor suppressor PALB2 (9). Notably, PALB2 does possess nucleosome binding activity and contacts the nucleosome acidic patch (37,38). Whether the nucleosome acidic patch also affects RAD51AP1 binding remains to be determined. Since most chromatin binding proteins - to ensure the specificity of their activity - seem to recognize multiple chromatin interfaces (38,39), we predict that the interaction between RAD51AP1 and chromatin is governed by multiple sites of contact and possibly other RAD51AP1 binding partners in cells.

## Material and methods

### Generation of the mono-Nucleosome Core Particle (NCP)

NCPs were reconstituted using a 147 bp dsDNA fragment with positioning sequence (601 Widom fragment) and human histone octamers using salt gradient deposition (12,14,24).

### Electrophoretic mobility shift assay

The NCP or the 147 bp dsDNA (40) (0.2 μM each) and purified full-length RAD51AP1, RAD54, RAD51AP1-F1, -F2, -F3, -N88, -C60 fragments or RAD51AP1-F3 mutants (0.1-0.8 μM) were incubated in buffer A (50 mM Tris-HCl pH 7.5, 100 mM NaCl) at 4°C for 30 min. Samples were fractionated by native 5% polyacrylamide gel electrophoresis (PAGE) in 0.2× TBE and 150 V for 1 h. Gels were stained by ethidium bromide (EtBr; Apex) or Imperial protein stain (ThermoScientific) and image acquisition and analyses were performed on a ChemiDoc XRS+ instrument equipped with Image Lab 6.1 software (BioRAD).

### Purification of RAD51AP1, RAD51 fragments, RAD51AP1 mutants and other proteins

The purification procedures for full-length human RAD51AP1, the RAD51AP1 fragments and mutants, and for human RAD54, RAD51 and NAP1 have been described elsewhere (5,6,11,24,41,42). Purified human histone octamer was purchased from The Histone Source at Colorado State University (https://www.histonesource.com/)and purified as described (43).

For purification of the RAD51AP1-N88 protein, *E. coli* Rosetta cells harboring the plasmid were grown and induced for protein expression as described earlier (42). RAD51AP1-N88 was purified by tandem purification using Ni-NTA resin (ThermoScientific) first, as described (42). Eluted protein was incubated with pre-equilibrated Strep-Tactin Superflow Plus (Qiagen) and gentle rotation at 4°C for 1 h in binding buffer containing 50 mM Tris-HCl, pH 7.5, 150 mM NaCl and 2 mM DTT. The resin was washed 4× in 3 bed volumes of binding buffer, bound protein was eluted with binding buffer containing 2.5 mM d-Desthiobiotin (Millipore) and dialyzed into dialysis buffer (50 mM Tris-HCl, pH 7.5, 300 mM NaCl, 0.05% Triton X-100, 2 mM DTT and 20% glycerol) at 4°C overnight before snap-freezing, and stored at −80°C.

### Mutant construction

The generation of the RAD51AP1 fragments F1, F2, F3 and C60 is described elsewhere (6). K4A, K4A-1, K4A-2, K3WA, K3WA-1, K3WA-2 mutants of RAD51AP1-F3, and K4A and K7WA (i.e., K4A with K3WA) mutants of the full-length RAD51AP1 protein were generated by Q5 Site-Directed Mutagenesis kit (New England Biolabs) following the instructions of the manufacturer and the primer pairs listed in Table 1.

**Table 1.**
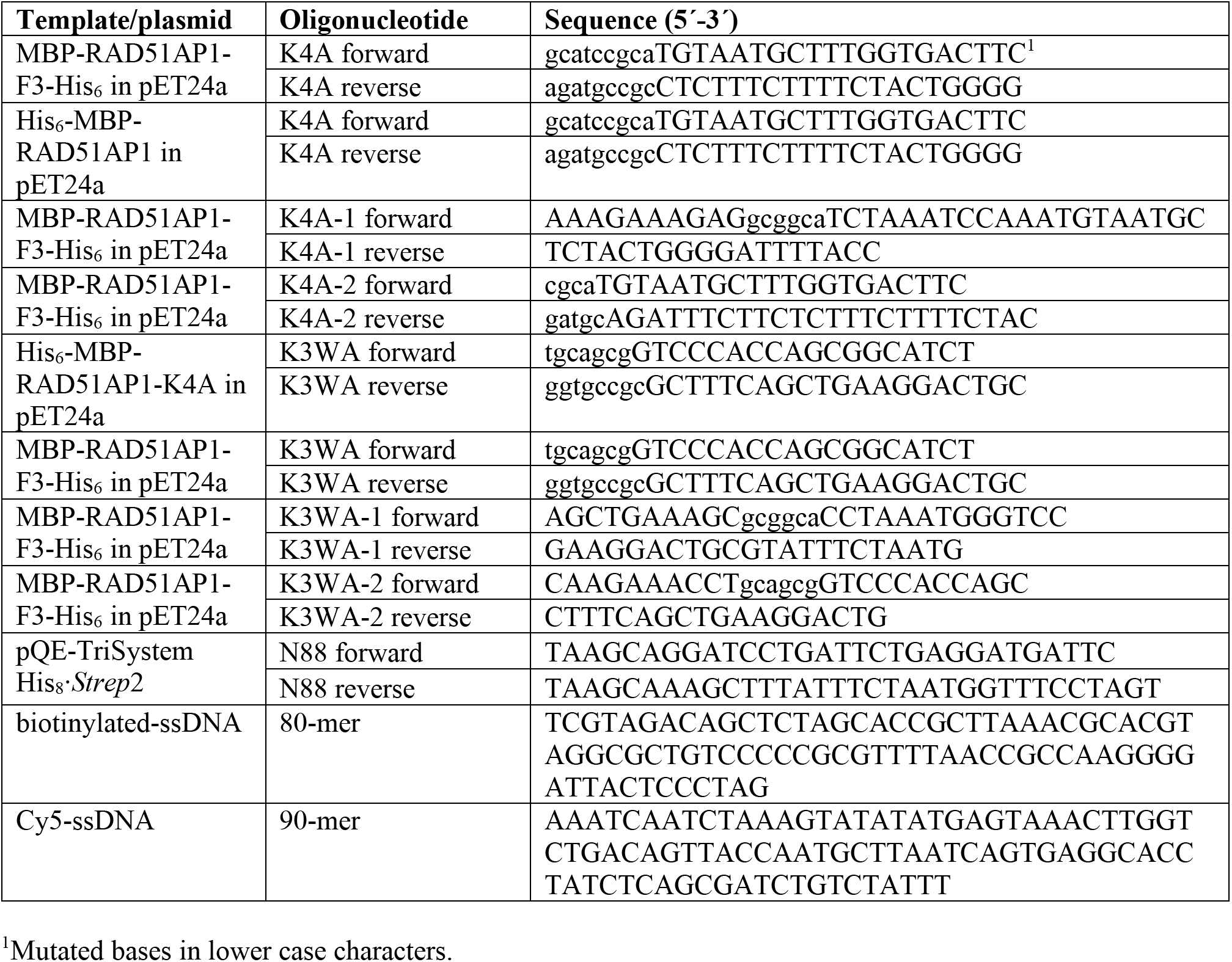
List of oligonucleotides/primers used in this study.

RAD51AP1-N88 was amplified from pOK24 (10) using the primer pairs listed in Table 1, and cloned from *Bam*HI to *Hind*III into pQE-TriSystem His·*Strep*2 (Qiagen) to be expressed with an N-terminal *Strep*-tag and a C-terminal His_8_-tag.

### Western blot analysis

Western blot analyses followed our standard protocols (11,44–46). The primary antibodies that were used are: α-RAD51AP1 (NB100-1129; Novus; 1:5,000; and our own α-RAD51AP1 antibody, as previously described in (9)), α-RAD54 (F-11; sc-374598; Santa Cruz Biotechnology; 1:500); α-RAD51 (Ab-1; EMD Millipore; 1:4,000), α-β-Actin (ab6276; Abcam; 1:3,000), α-H3 (ab1791; Abcam; 1:10,000), α-H2A (GTX1129418; GeneTex; 1:1,000); α-FLAG (F3165; Sigma; 1:1,000);α-MBP (PAI-989; ThermoScientific; 1:5,000). HRPconjugated goat anti-rabbit or goat anti-mouse IgG (Jackson ImmunoResearch Laboratories; 1:10,000) were used as secondary antibodies and SuperSignal Substrate kit (ThermoScientific) for the detection of signal.

### FAR Western analysis

Following the EMSA, proteins were electro-transferred onto a PVDF membrane. The membrane was first soaked for 2 h in buffer I (10 mM KH_2_PO_4_, pH 7.4, 150 mM KCl, 15 mg/ml BSA, 2 mM 2-mercaptoethanol, 0.05% Tween-20) prior to incubation with 3 μg/ml purified RAD51 in buffer I and at 4°C overnight. On the following day, the membrane was washed 3× in 10 ml buffer I, before incubation with primary α-RAD51 and secondary antibody (1 h each) in buffer I at room temperature and further detection of bound RAD51 protein by SuperSignal Substrate kit (ThermoScientific).

### Histone chaperone assay

The histone chaperone assay was performed essentially as described (18). Briefly, 0.5 μM H2A/H2B and 0.5 μM H3/H4 were pre-incubated with increasing amounts of RAD51AP1 (1.4-22.4 μM) or NAP1 (1.2 and 5.0 μM) in buffer D (25 mM Tris-HCl, pH 7.5, 100 mM NaCl, 1.5 mM MgCl_2_ and 1 mM DTT) at 37°C for 15 min. Then, the 147 bp DNA fragment containing the nucleosome positioning sequence (40) was added and samples were incubated further at 37°C for 15 min. The reaction was stopped by the addition of 20% sucrose to a final concentration of 5%, and samples were separated on a native 5% PAGE in 0.2× TBE buffer for 1 h at 150V. DNA was visualized by EtBr staining and image acquisition was performed on a ChemiDoc XRS+ instrument equipped with Image Lab 6.1 software (BioRAD).

### Size exclusion chromatography (SEC) – micro-HPLC

The NCP (2.5 μM) and MBP-RAD51AP1-F3 (2.5 μM) were incubated in buffer E (50 mM Tris-HCl, pH 7.5, 100 mM NaCl) in a final volume of 50 μl at 4°C for 15 min. Thirty μl of this reaction was injected into a 2.4 ml prepacked analytical Superdex-200 Increase 3.2/300 gel filtration column (GE Healthcare) pre-equilibrated with buffer E. Gel filtration standard (1.35-670 kD, pI 4.5-6.9; BioRad) was used to ensure that the column was properly packed and the samples were evenly eluted. Chromatography was conducted at a flow rate of 0.075 ml/min and absorbance was monitored at 280 nm. Fractions of 0.1 ml were collected and analyzed by agarose gel electrophoresis and Western blots for the presence of histone H3, DNA and RAD51AP1-F3 to confirm co-elution of the latter with the NCP.

### In-vitro affinity pull-down assays

MBP- and His_6_-tagged full-length RAD51AP1 or MBP- and His_6_-tagged RAD51AP1-F1, -F2, -F3 (80 nM each) were incubated with pre-equilibrated Ni-NTA resin (Thermofisher Scientific) in buffer F (50 mM Tris-HCl, pH 7.5, 150 mM NaCl, 0.1% Triton X-100, 2% BSA) at 4°C with gentle rotation for 1 h. The supernatant was removed and bound protein was further incubated with histone Octamer (80 nM) in buffer F containing DNase (1 U/μg protein) and at 4°C with gentle rotation for 2 h. The resin then was washed 4× with buffer F and bound protein complexes were eluted in 40 μl buffer F with 300 mM imidazole. Eluted complexes (10 μl) were fractionated on a 10% NuPAGE protein gel (Thermofisher Scientific), transferred onto a PVDF membrane and detected by Western blot analysis.

### Duplex capture assay

The duplex capture assay was performed essentially as previously described (47). Briefly, the presynaptic filament was formed by incubating 5 μl of streptavidin-coated magnetic resin (Roche Molecular Biochemicals) with 5’-biotinylated 80-mer ssDNA oligonucleotide (5 μM; Table 1) and RAD51 (700 nM) in buffer G (25 mM Hepes, pH 7.5, 50 mM Tris-HCl, pH 7.5, 35 mM NaCl, 45 mM KCl, 1 mM MgCl_2_, 0.16 mM EDTA, 2 mM ATP, 2% glycerol, 0.01% NP40, 0.4 mM β-mercaptoethanol, and 100 μg/ml BSA) at 37°C for 5 min. The resin was captured magnetically and washed once with 20 μl buffer G. The wash was removed and resin was resuspended in 10 μl buffer D containing full-length FLAG-tagged RAD51AP1 (400 or 800 nM). After a 5-min incubation at 37°C, the resin was captured and washed as before. The supernatant was removed and 10 μl buffer G containing either the 147 bp dsDNA or the NCP (1 μM each) were added and incubated at 37°C for 10 min. The resin was captured, the supernatant was saved, and the resin was washed 4× with 200 μl buffer G. Both resin and supernatant were treated with 2 mg/ml ProteinaseK in ProteinaseK buffer (2 mM Tris-HCl, pH 7.5, 1 mM CaCl_2_, 0.2% SDS) at 37°C for 15 min, and supernatant-containing and resin-bound DNA were analyzed by 1% agarose gel electrophoresis. DNA was visualized by EtBr staining and image acquisition was performed on a ChemiDoc XRS+ instrument equipped with Image Lab 6.1 software (BioRAD). Quantification of signal intensities was done by ImageJ (https://imagej.nih.gov/).

### Chromatin assembly

Chromatin was assembled with human histone octamer by salt gradient dialysis on pBluescript II SK(-) plasmid DNA, as described (48). The plasmid/octamer ratio was based on 207 ± 4 bp DNA/nucleosome. The quality of the assembled chromatin was controlled by limited digestion with MNase.

### D-loop assay

The D-loop assay essentially was performed in 50 mM KCl and as described earlier (11,42,49), except that Cy5-labeled 90-mer ssDNA purchased from Integrated DNA Technologies (IDT) was used (Table 1). The 90-mer is homologous to the region of 1932-2022 bp within the pBluescript II SK(-) plasmid DNA (50). Fluorescence image capture and analysis was conducted on an Odyssey CLx Imaging System equipped with Image Studio 5.0 software (LI-COR).

### Cell culture, cell fractionation and treatment with mitomycin C

U2OS and HT1080 cells were obtained from ATCC and maintained as recommended. Cell fractionation was carried out using the Subcellular Protein Fractionation Kit (ThermoFisher Scientific) and as described by the manufacturer. Exposure of cells to 1 μM mitomycin C (MMC; Sigma) occurred in regular growth medium at 37°C for 24 h.

### Statistical analyses

Statistical analyses were performed using Prism 8 GraphPad Software on the data from 2-4 independent experiments. Statistical significance was assessed by two-way ANOVA or multiple *t*-test analysis, as indicated. *P* ≤ 0.05 was considered significant.

## Data availability

All data are included in the manuscript and in the supporting information. The raw data are available upon request.

## Author contributions

N.S. and C.W. conceptualization; E.P., N.S., P.S., B.W., Y.H. and W.Z. data curation; E.P., N.S., B.W. and W.Z. data analyses and validation; D.S.A. provided reagents; E.P., N.S. and C.W. writing of the manuscript; W.Z. and C.W. funding acquisition.

## Funding and additional information

This work was supported by National Institutes of Health Grants R01ES021454, R56ES021454, R03ES029206 (to C.W.), a V Foundation Scholar Grant and a Max & Minnie Tomerlin Voelcker Young Investigator Award (to W.Z.). E.P. was co-supported by T32OD012201. The content is solely the responsibility of the authors and does not necessarily represent the official views of the National Institutes of Health.

## Acknowledgements

*We thank Drs. Karolin Luger and Uma M. Muthurajan for materials, guidance and comments on the manuscript*.

## Abbreviations

The abbreviations used are:

RAD51AP1: RAD51 Associated Protein 1
HR: Homologous Recombination
NCP: Nucleosome Core Particle
D-loop: Displacement-loop
EMSA: Electrophoretic Mobility Shift Assay
AA: amino acid
MMC: mitomycin C
PVDF: Polyvinylidene Difluoride
SEC: size exclusion chromatography
HPLC: high pressure liquid chromatography
Ni-NTA: nickel nitrilotriacetic acid
BSA: Bovine Serum Albumin
MBP: Maltose Binding Protein
NAP1: Nucleosome-Associated Protein 1
Hop2: Homologous pairing 2
Mnd1: Meiotic nuclear divisions 1
PALB2: Partner and Localizer of BRCA2
UAF1: USP1-Associated Factor 1
ATP: Adenosine Triphosphate
MNase: micrococcal nuclease
ANOVA: analysis of variance.

## References

1. Tye, S., Ronson, G. E., and Morris, J. R. (2020) A fork in the road: Where homologous recombination and stalled replication fork protection part ways. Semin Cell Dev Biol

2. Rambo, R. P., and Tainer, J. A. (2013) Accurate assessment of mass, models and resolution by small-angle scattering. Nature 496, 477–481

3. Kulkarni, P., and Uversky, V. N. (2018) Intrinsically Disordered Proteins: The Dark Horse of the Dark Proteome. Proteomics 18, e1800061

4. Modesti, M., Budzowska, M., Baldeyron, C., Demmers, J. A., Ghirlando, R., and Kanaar, R. (2007) RAD51AP1 is a structure-specific DNA binding protein that stimulates joint molecule formation during RAD51-mediated homologous recombination. Molecular cell 28, 468–481

5. Wiese, C., Dray, E., Groesser, T., San Filippo, J., Shi, I., Collins, D. W., Tsai, M. S., Williams, G. J., Rydberg, B., Sung, P., and Schild, D. (2007) Promotion of homologous recombination and genomic stability by RAD51AP1 via RAD51 recombinase enhancement. Molecular cell 28, 482–490

6. Dunlop, M. H., Dray, E., Zhao, W., San Filippo, J., Tsai, M. S., Leung, S. G., Schild, D., Wiese, C., and Sung, P. (2012) Mechanistic insights into RAD51-associated protein 1 (RAD51AP1) action in homologous DNA repair. The Journal of biological chemistry 287, 12343–12347

7. Dunlop, M. H., Dray, E., Zhao, W., Tsai, M. S., Wiese, C., Schild, D., and Sung, P. (2011) RAD51-associated protein 1 (RAD51AP1) interacts with the meiotic recombinase DMC1 through a conserved motif. J Biol Chem 286, 37328–37334

8. Dray, E., Dunlop, M. H., Kauppi, L., San Filippo, J., Wiese, C., Tsai, M. S., Begovic, S., Schild, D., Jasin, M., Keeney, S., and Sung, P. (2011) Molecular basis for enhancement of the meiotic DMC1 recombinase by RAD51 associated protein 1 (RAD51AP1). Proc Natl Acad Sci U S A 108, 3560–3565

9. Dray, E., Etchin, J., Wiese, C., Saro, D., Williams, G. J., Hammel, M., Yu, X., Galkin, V. E., Liu, D., Tsai, M. S., Sy, S. M., Schild, D., Egelman, E., Chen, J., and Sung, P. (2010) Enhancement of RAD51 recombinase activity by the tumor suppressor PALB2. Nat Struct Mol Biol 17, 1255–1259

10. Kovalenko, O. V., Golub, E. I., Bray-Ward, P., Ward, D. C., and Radding, C. M. (1997) A novel nucleic acid-binding protein that interacts with human rad51 recombinase. Nucleic Acids Res 25, 4946–4953

11. Parplys, A. C., Zhao, W., Sharma, N., Groesser, T., Liang, F., Maranon, D. G., Leung, S. G., Grundt, K., Dray, E., Idate, R., Ostvold, A. C., Schild, D., Sung, P., and Wiese, C. (2015) NUCKS1 is a novel RAD51AP1 paralog important for homologous recombination and genome stability. Nucleic Acids Res 43, 9817–9834

12. Gottesfeld, J. M., and Luger, K. (2001) Energetics and affinity of the histone octamer for defined DNA sequences. Biochemistry 40, 10927–10933

13. Lusser, A., and Kadonaga, J. T. (2004) Strategies for the reconstitution of chromatin. Nat Methods 1, 19–26

14. Dyer, P. N., Edayathumangalam, R. S., White, C. L., Bao, Y., Chakravarthy, S., Muthurajan, U. M., and Luger, K. (2004) Reconstitution of nucleosome core particles from recombinant histones and DNA. Methods Enzymol 375, 23–44

15. Alexeev, A., Mazin, A., and Kowalczykowski, S. C. (2003) Rad54 protein possesses chromatin-remodeling activity stimulated by the Rad51-ssDNA nucleoprotein filament. Nat Struct Biol 10, 182–186

16. Zhang, Z., Fan, H. Y., Goldman, J. A., and Kingston, R. E. (2007) Homology-driven chromatin remodeling by human RAD54. Nat Struct Mol Biol 14, 397–405

17. Hammond, C. M., Stromme, C. B., Huang, H., Patel, D. J., and Groth, A. (2017) Histone chaperone networks shaping chromatin function. Nat Rev Mol Cell Biol 18, 141–158

18. Sato, K., Ishiai, M., Toda, K., Furukoshi, S., Osakabe, A., Tachiwana, H., Takizawa, Y., Kagawa, W., Kitao, H., Dohmae, N., Obuse, C., Kimura, H., Takata, M., and Kurumizaka, H. (2012) Histone chaperone activity of Fanconi anemia proteins, FANCD2 and FANCI, is required for DNA crosslink repair. EMBO J 31, 3524–3536

19. Higgs, M. R., Sato, K., Reynolds, J. J., Begum, S., Bayley, R., Goula, A., Vernet, A., Paquin, K. L., Skalnik, D. G., Kobayashi, W., Takata, M., Howlett, N. G., Kurumizaka, H., Kimura, H., and Stewart, G. S. (2018) Histone Methylation by SETD1A Protects Nascent DNA through the Nucleosome Chaperone Activity of FANCD2. Molecular cell 71, 25–41 e26

20. Mehrotra, P. V., Ahel, D., Ryan, D. P., Weston, R., Wiechens, N., Kraehenbuehl, R., Owen-Hughes, T., and Ahel, I. (2011) DNA repair factor APLF is a histone chaperone. Molecular cell 41, 46–55

21. Muthurajan, U. M., Hepler, M. R., Hieb, A. R., Clark, N. J., Kramer, M., Yao, T., and Luger, K. (2014) Automodification switches PARP-1 function from chromatin architectural protein to histone chaperone. Proc Natl Acad Sci U S A 111, 12752–12757

22. Fujii-Nakata, T., Ishimi, Y., Okuda, A., and Kikuchi, A. (1992) Functional analysis of nucleosome assembly protein, NAP-1. The negatively charged COOH-terminal region is not necessary for the intrinsic assembly activity. J Biol Chem 267, 20980–20986

23. Park, Y. J., and Luger, K. (2006) The structure of nucleosome assembly protein 1. Proc Natl Acad Sci U S A 103, 1248–1253

24. Sharma, N., and Nyborg, J. K. (2008) The coactivators CBP/p300 and the histone chaperone NAP1 promote transcription-independent nucleosome eviction at the HTLV-1 promoter. Proc Natl Acad Sci U S A 105, 7959–7963

25. Kovalenko, O. V., Wiese, C., and Schild, D. (2006) RAD51AP2, a novel vertebrate- and meiotic-specific protein, shares a conserved RAD51-interacting C-terminal domain with RAD51AP1/PIR51. Nucleic Acids Res 34, 5081–5092

26. Liang, F., Longerich, S., Miller, A. S., Tang, C., Buzovetsky, O., Xiong, Y., Maranon, D. G., Wiese, C., Kupfer, G. M., and Sung, P. (2016) Promotion of RAD51-Mediated Homologous DNA Pairing by the RAD51AP1-UAF1 Complex. Cell reports 15, 2118–2126

27. Liang, F., Miller, A. S., Tang, C., Maranon, D., Williamson, E. A., Hromas, R., Wiese, C., Zhao, W., Sung, P., and Kupfer, G. M. (2020) The DNA-binding activity of USP1-associated factor 1 is required for efficient RAD51-mediated homologous DNA pairing and homology-directed DNA repair. J Biol Chem 295, 8186–8194

28. Raynard, S., and Sung, P. (2009) Assay for human Rad51-mediated DNA displacement loop formation. Cold Spring Harb Protoc 2009, pdb prot5120

29. Jaskelioff, M., Van Komen, S., Krebs, J. E., Sung, P., and Peterson, C. L. (2003) Rad54p is a chromatin remodeling enzyme required for heteroduplex DNA joint formation with chromatin. J Biol Chem 278, 9212–9218

30. Alexiadis, V., and Kadonaga, J. T. (2002) Strand pairing by Rad54 and Rad51 is enhanced by chromatin. Genes & development 16, 2767–2771

31. Kwon, Y., Seong, C., Chi, P., Greene, E. C., Klein, H., and Sung, P. (2008) ATP-dependent chromatin remodeling by the Saccharomyces cerevisiae homologous recombination factor Rdh54. J Biol Chem 283, 10445–10452

32. Daley, J. M., Gaines, W. A., Kwon, Y., and Sung, P. (2014) Regulation of DNA pairing in homologous recombination. Cold Spring Harb Perspect Biol 6, a017954

33. Wright, W. D., Shah, S. S., and Heyer, W. D. (2018) Homologous recombination and the repair of DNA double-strand breaks. J Biol Chem 293, 10524–10535

34. Pires, E., Sung, P., and Wiese, C. (2017) Role of RAD51AP1 in homologous recombination DNA repair and carcinogenesis. DNA repair 59, 76–81

35. Crickard, J. B., Kwon, Y., Sung, P., and Greene, E. C. (2019) Dynamic interactions of the homologous pairing 2 (Hop2)-meiotic nuclear divisions 1 (Mnd1) protein complex with meiotic presynaptic filaments in budding yeast. J Biol Chem 294, 490–501

36. Liang, F., Miller, A. S., Longerich, S., Tang, C., Maranon, D., Williamson, E. A., Hromas, R., Wiese, C., Kupfer, G. M., and Sung, P. (2019) DNA requirement in FANCD2 deubiquitination by USP1-UAF1-RAD51AP1 in the Fanconi anemia DNA damage response. Nat Commun 10, 2849

37. Bleuyard, J. Y., Buisson, R., Masson, J. Y., and Esashi, F. (2012) ChAM, a novel motif that mediates PALB2 intrinsic chromatin binding and facilitates DNA repair. EMBO Rep 13, 135–141

38. Belotserkovskaya, R., Raga Gil, E., Lawrence, N., Butler, R., Clifford, G., Wilson, M. D., and Jackson, S. P. (2020) PALB2 chromatin recruitment restores homologous recombination in BRCA1-deficient cells depleted of 53BP1. Nat Commun 11, 819

39. Wilson, M. D., and Durocher, D. (2017) Reading chromatin signatures after DNA double-strand breaks. Philos Trans R Soc Lond B Biol Sci 372

40. Lowary, P. T., and Widom, J. (1998) New DNA sequence rules for high affinity binding to histone octamer and sequence-directed nucleosome positioning. J Mol Biol 276, 19–42

41. Sigurdsson, S., Van Komen, S., Bussen, W., Schild, D., Albala, J. S., and Sung, P. (2001) Mediator function of the human Rad51B-Rad51C complex in Rad51/RPA-catalyzed DNA strand exchange. Genes & development 15, 3308–3318

42. Maranon, D. G., Sharma, N., Huang, Y., Selemenakis, P., Wang, M., Altina, N., Zhao, W., and Wiese, C. (2020) NUCKS1 promotes RAD54 activity in homologous recombination DNA repair. J Cell Biol 219

43. Strickfaden, H., Tolsma, T. O., Sharma, A., Underhill, D. A., Hansen, J. C., and Hendzel, M. J. (2020) Condensed Chromatin Behaves like a Solid on the Mesoscale In Vitro and in Living Cells. Cell 183, 1772–1784 e1713

44. Wiese, C., Hinz, J. M., Tebbs, R. S., Nham, P. B., Urbin, S. S., Collins, D. W., Thompson, L. H., and Schild, D. (2006) Disparate requirements for the Walker A and B ATPase motifs of human RAD51D in homologous recombination. Nucleic Acids Res 34, 2833–2843

45. Parplys, A. C., Kratz, K., Speed, M. C., Leung, S. G., Schild, D., and Wiese, C. (2014) RAD51AP1-deficiency in vertebrate cells impairs DNA replication. DNA repair 24, 87–97

46. Zhao, W., Vaithiyalingam, S., San Filippo, J., Maranon, D. G., Jimenez-Sainz, J., Fontenay, G. V., Kwon, Y., Leung, S. G., Lu, L., Jensen, R. B., Chazin, W. J., Wiese, C., and Sung, P. (2015) Promotion of BRCA2-Dependent Homologous Recombination by DSS1 via RPA Targeting and DNA Mimicry. Molecular cell 59, 176–187

47. Kobayashi, W., Takaku, M., Machida, S., Tachiwana, H., Maehara, K., Ohkawa, Y., and Kurumizaka, H. (2016) Chromatin architecture may dictate the target site for DMC1, but not for RAD51, during homologous pairing. Scientific reports 6, 24228

48. Jeong, S. W., Lauderdale, J. D., and Stein, A. (1991) Chromatin assembly on plasmid DNA in vitro. Apparent spreading of nucleosome alignment from one region of pBR327 by histone H5. J Mol Biol 222, 1131–1147

49. Kwon, Y., Chi, P., Roh, D. H., Klein, H., and Sung, P. (2007) Synergistic action of the Saccharomyces cerevisiae homologous recombination factors Rad54 and Rad51 in chromatin remodeling. DNA repair 6, 1496–1506

50. Kwon, Y., Daley, J. M., and Sung, P. (2017) Reconstituted System for the Examination of Repair DNA Synthesis in Homologous Recombination. Methods Enzymol 591, 307–325

