## Supporting Information - Figure S1-S4 for "RAD51AP1 mediates RAD51 activity through nucleosome interaction"

**Supporting Materials and methods**

**Supporting Figure S1**

**Supporting Figure S2**

**Supporting Figure S3**

**Supporting Figure S4**

**Supporting References**

### Supporting Materials and methods

*Determination of binding constants*

Binding constants were determined by EMSA and ethidium bromide staining with purified $His_6$-/FLAG-tagged full-length RAD51AP1 (8-270 nM) and the NCP (0.2 µM) or the Widom 601 147 bp dsDNA fragment (0.1 µM). Integrated densitometry values (IDVs) of the non-shifted bands (ns) were assessed by Image Lab 6.1 software (BioRad) and the fraction of protein bound (Y) was determined as $Y = 1 - (IDV_{ns} - IDV_{background})$. GraphPad Prism9 software and nonlinear regression were used to obtain apparent binding constants ($K_{D(app)}$). To account for the diminished efficiency of ethidium bromide in staining nucleosome-associated DNA, a normalization factor of 5.2 was determined based on 3 independent native gels comparing the signal intensities of ethidium bromide for different amounts of NCP to the 147 bp dsDNA fragment (see Fig. S2*J* and *K*).

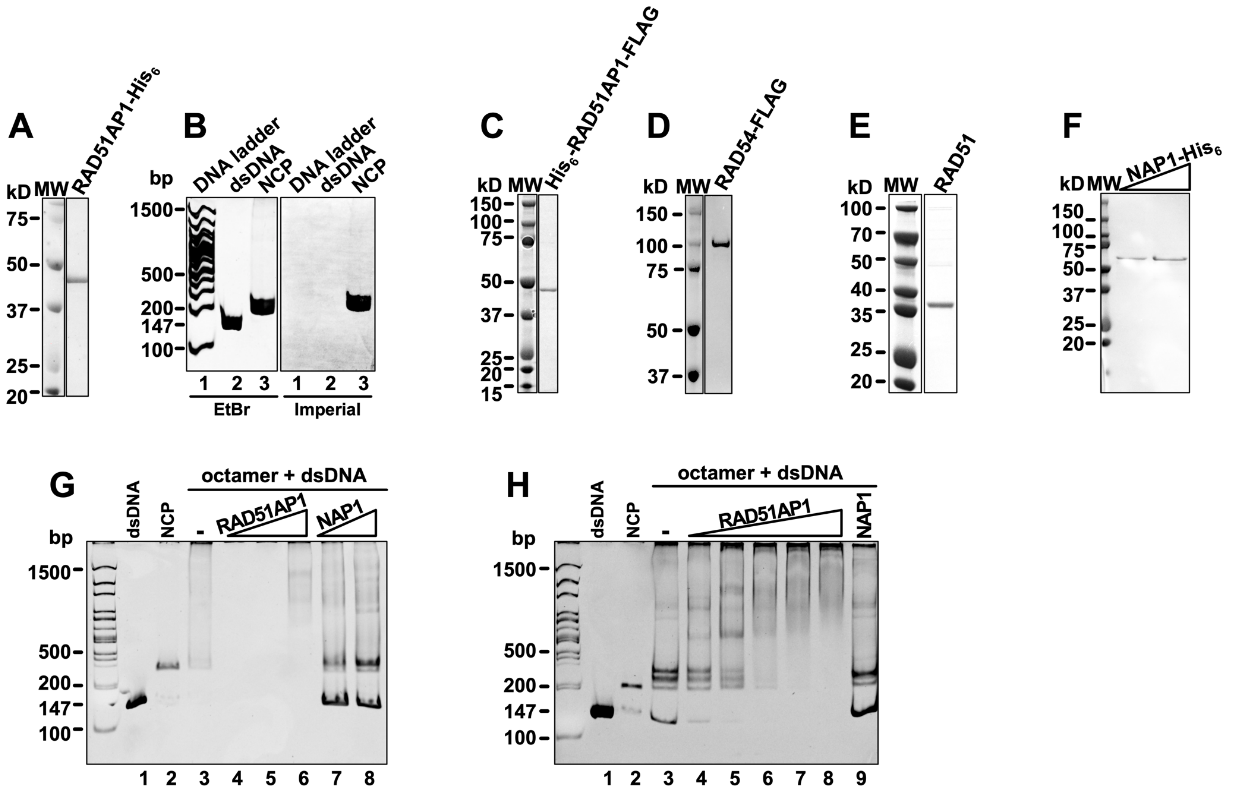


**Figure S1. RAD51AP1 does not have histone chaperone activity.** ***A***, SDS-PAGE of purified RAD51AP1-His$_6$. ***B***, Imperial protein stain does not stain DNA. EMSA of 147 bp dsDNA and NCP (lane 2 and 3, respectively) stained by ethidium bromide (EtBr) first and then by Imperial protein stain. ***C-F***, SDS-PAGE of purified proteins (His$_6$-RAD51AP1-FLAG, RAD54-FLAG, RAD51 and NAP1-His$_6$, respectively). ***G-H***, RAD51AP1 does not have histone chaperone activity. ***G***, EMSA of dsDNA (147 bp) with H2A/H2B and H3/H4 (DNA:histones (1:1)) and 1.4, 2.8 and 5.6 μM His$_6$-/FLAG-tagged full-length RAD51AP1 (lanes 4-6) or 2.5 and 5 μM His$_6$-tagged NAP1 (lanes 7 and 8). ***H***, EMSA of dsDNA (147 bp) with H2A/H2B and H3/H4 (dsDNA:histones (1:1)) and 2.8, 5.6, 11.2, 16.8 and 22.4 μM His$_6$-/FLAG-tagged RAD51AP1 (lanes 4-8) and 1.2 μM His$_6$-tagged NAP1 (lane 9).

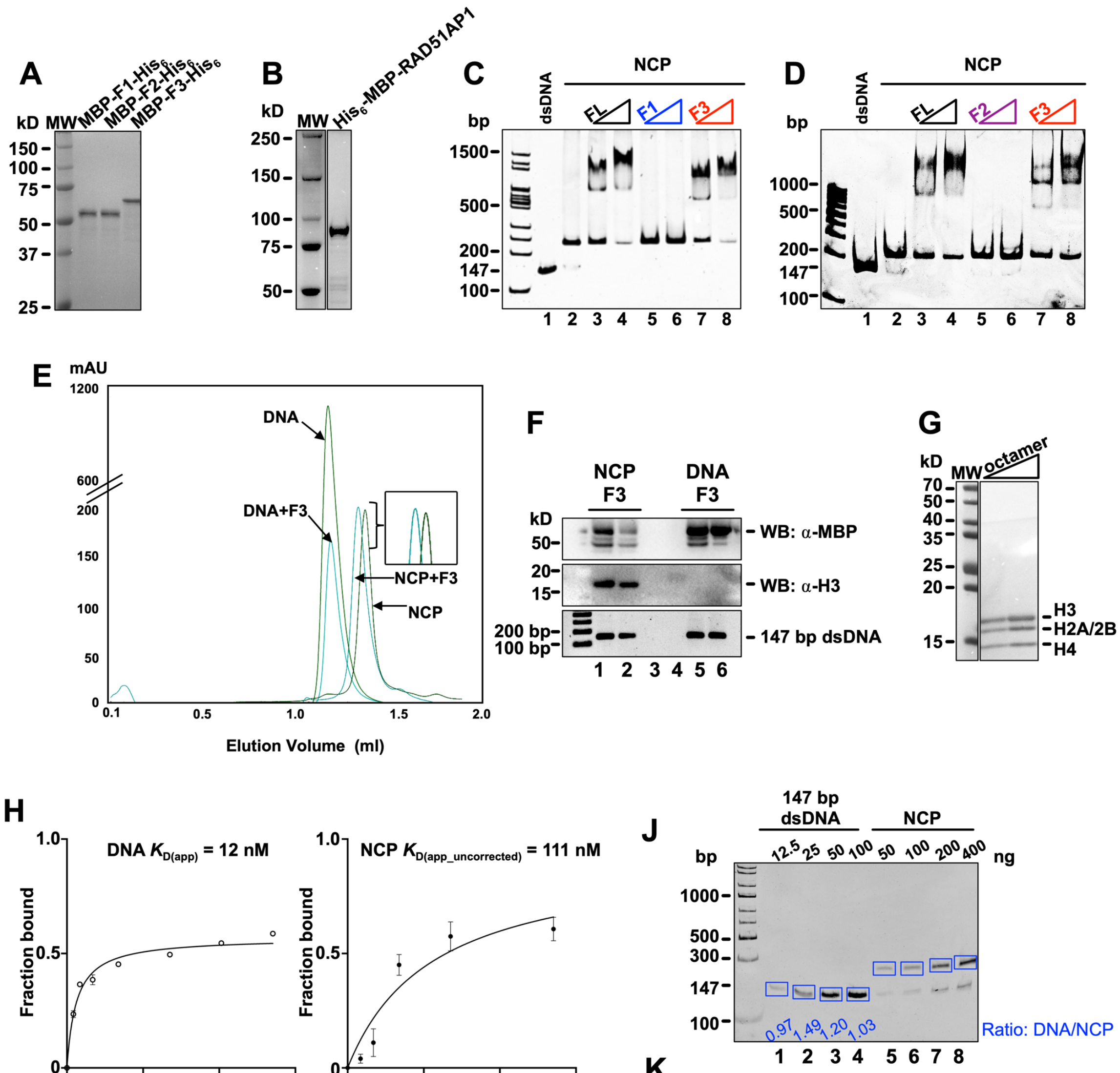


K

| | Lane | Net Band | | | DNA/NCP sum | | | Mean normalized | x 4 (ng loaded) | AVERAGE |
|---|---|---|---|---|---|---|---|---|---|---|
| | | Gel 1 | Gel 2 | Gel 3 | Gel 1 | Gel 2 | Gel 3 | | | |
| DNA | 1 | 17.096 | 11.047 | 3.483 | | | | | | |
| | 2 | 35.464 | 20.24 | 10.922 | | | | | | |
| | 3 | 60.945 | 38.764 | 25.274 | | | | | | |
| | 4 | 66.439 | 52.803 | 32.228 | | | | | | |
| NCP sum | 5 | 8.918 | 9.668 | 3.597 | 1.917022 | 1.142635 | 0.968307 | 1.34 | 5.37 | 5.20 |
| | 6 | 25.729 | 16.258 | 7.327 | 1.378367 | 1.244926 | 1.490651 | 1.37 | 5.49 | |
| | 7 | 56.448 | 30.471 | 21.014 | 1.079666 | 1.27216 | 1.202722 | 1.18 | 4.74 | |
| | 8 | 81.787 | 92.122 | 31.189 | 0.812342 | 0.573186 | 1.033313 | 0.81 | 3.23 | |

→ NCP $K_{D\,(app\_corrected)}$ = 111/5.2 = 21 nM

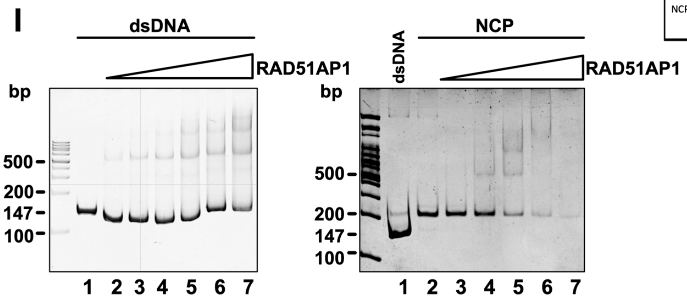

**Figure S2. The RAD51AP1-F3 fragment interacts with the NCP.** ***A,*** SDS-PAGE of purified MBP/$His_6$-tagged RAD51AP1-F1, -F2 and -F3 fragments. ***B,*** SDS-PAGE of purified MBP/$His_6$-tagged full-length RAD51AP1. ***C,*** The RAD51AP1-F1 fragment does not bind to the NCP (lanes 5 and 6). EMSA of full-length MBP/$His_6$-RAD51AP1 (here: FL), F1 or F3 (0.1 and 0.2 µM each) with the NCP (0.2 µM). ***D,*** The RAD51AP1-F2 fragment does not bind to the NCP (lanes 5 and 6). EMSA of FL MBP/$His_6$-RAD51AP1, F2 or F3 (0.1 and 0.2 µM each) with the NCP (0.2 µM). ***E***, NCP/F3 and dsDNA/F3 complexes co-elute from size exclusion chromatography (SEC). Overlay of the chromatograms. ***F***, Western blots and agarose gel of the eluted fractions: Co-elution of RAD51AP1-F3, histone H3 and the 147 bp dsDNA (from NCP) (lanes 1 and 2), and co-elution of RAD51AP1-F3 and the 147 bp dsDNA (lanes 5 and 6). ***G,*** SDS-PAGE of purified human histone octamer. ***H***, Determination of the apparent binding constants ($K_{D(app)}$) of $His_6$-/FLAG-RAD51AP1 for the Widom 601 147 bp dsDNA fragment (left panel) or the NCP (right panel). Data points are the means from 3 independent experiments ± 1SD. $K_{D(app)}$ (dsDNA) = 12 nM (d.f. = 4; $r^2$ = 0.91) and the $K_{D(app/uncorrected)}$ (NCP) = 111 nM (d.f. = 6; $r^2$ = 0.98). ***I***, Representative EMSAs of $His_6$-/FLAG-RAD51AP1 with the Wisom 601 147 bp dsDNA fragment (left panel; 8-270 nM RAD51AP1) and the NCP (right panel; 16-270 nM RAD51AP1) stained by ethidium bromide (EtBr) for the determination of binding constants in *H*. ***J***, Representative native PAGE stained with EtBr and used for densitometry to assess difference in staining efficiency between dsDNA and nucleosomal dsDNA. ***K***, Calculation of normalization factor for EtBr between dsDNA and nucleosomal dsDNA from three independent gels and correction of $K_{D(app/corrected)}$ (NCP) = 111/5.2 nM to 21 nM.

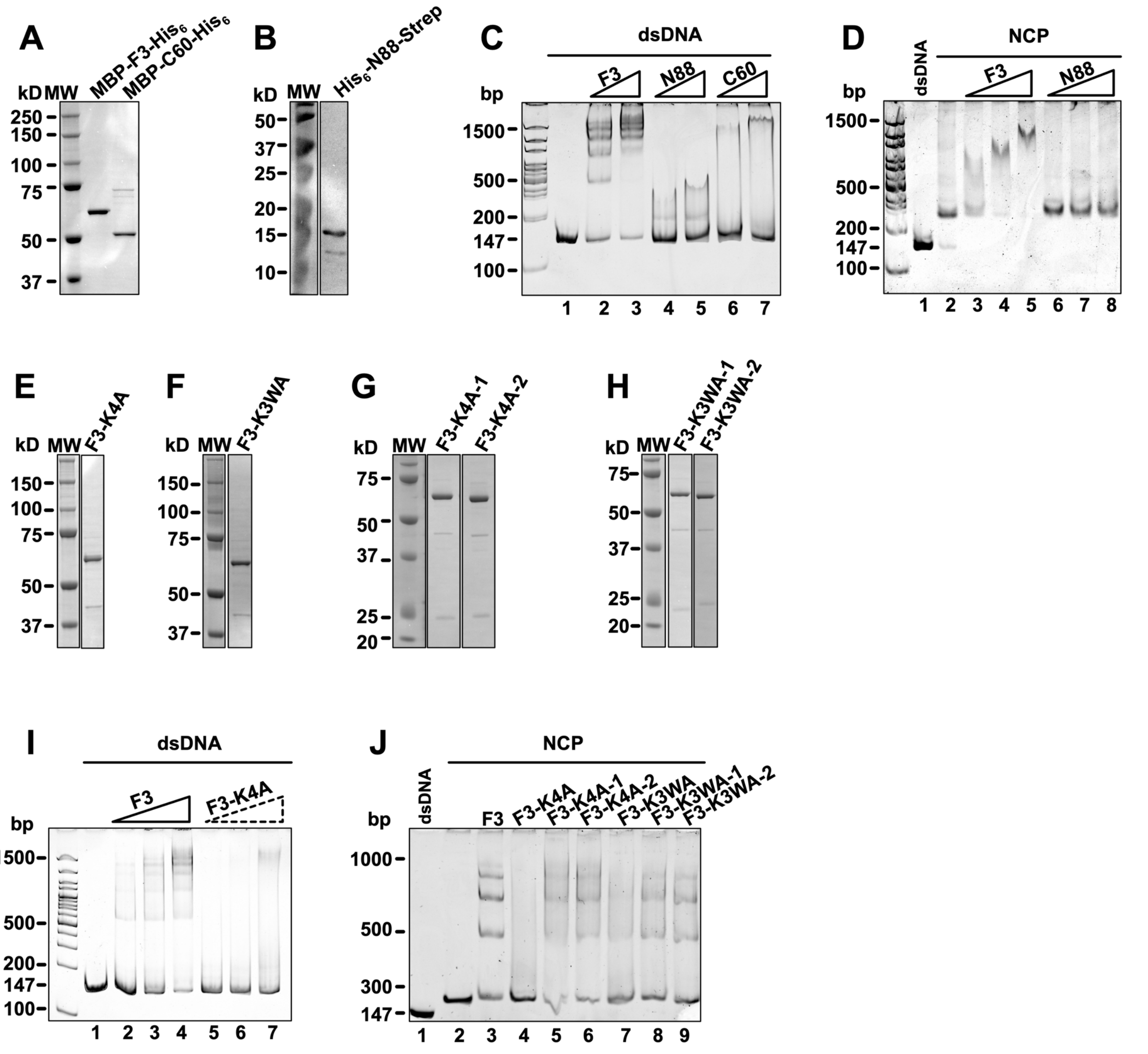


**Figure S3. Deletion or mutation of the DNA binding domain negatively affects the ability of RAD51AP1-F3 to associate with the NCP.** ***A-B***, SDS-PAGE of the purified RAD51AP1-F3 fragments: F3, F3-C60 and F3-N88. ***C***, F3-N88 and F3-C60 are greatly impaired in DNA binding. EMSA of RAD51AP1-F3, -N88 and -C60 (0.2 and 0.4 μM each) with 147 bp dsDNA (0.2 μM). ***D,*** F3-N88 does not bind to the NCP. EMSA of RAD51AP1-F3 and -N88 (0.2, 0.4 and 0.8 μM each) with the NCP (0.2 μM). ***E-H***, SDS-PAGE of purified RAD51AP1-F3 mutants. ***I***, F3-K4A (compound point mutant of *K231A, K232A, K234A and K236A) does not bind to dsDNA (147 bp), as previously shown (1). EMSA of RAD51AP1-F3 and RAD51AP1-F3-K4A (0.25, 0.5 and 1.0 μM each) with dsDNA (0.2 μM). ***J***, DNA binding mutants of RAD51AP1-F3 (0.3 μM each) are defective (F3-K4A; lane 4) or greatly impaired (F3-K4A-1, -K4A-2, -K3WA, K3WA-1 and K3WA-2; lanes 5-9) in their association with the NCP (0.2 μM). RAD51AP1-F3 (0.3 μM; lane 3) is shown for comparison. F3-KA4-1: compound point mutant of K231A and K232A; F3-KA4-2: compound point mutant of K234A and K236A; F3-K3WA: compound point mutant of K283A, K284A, K286A and W287A; F3-K3WA-1: compound point mutant of K283A and K284A; F3-K3WA-2: compound point mutant of K286A and W287A. *Residue numbering based on RAD51AP1 isoform 2.

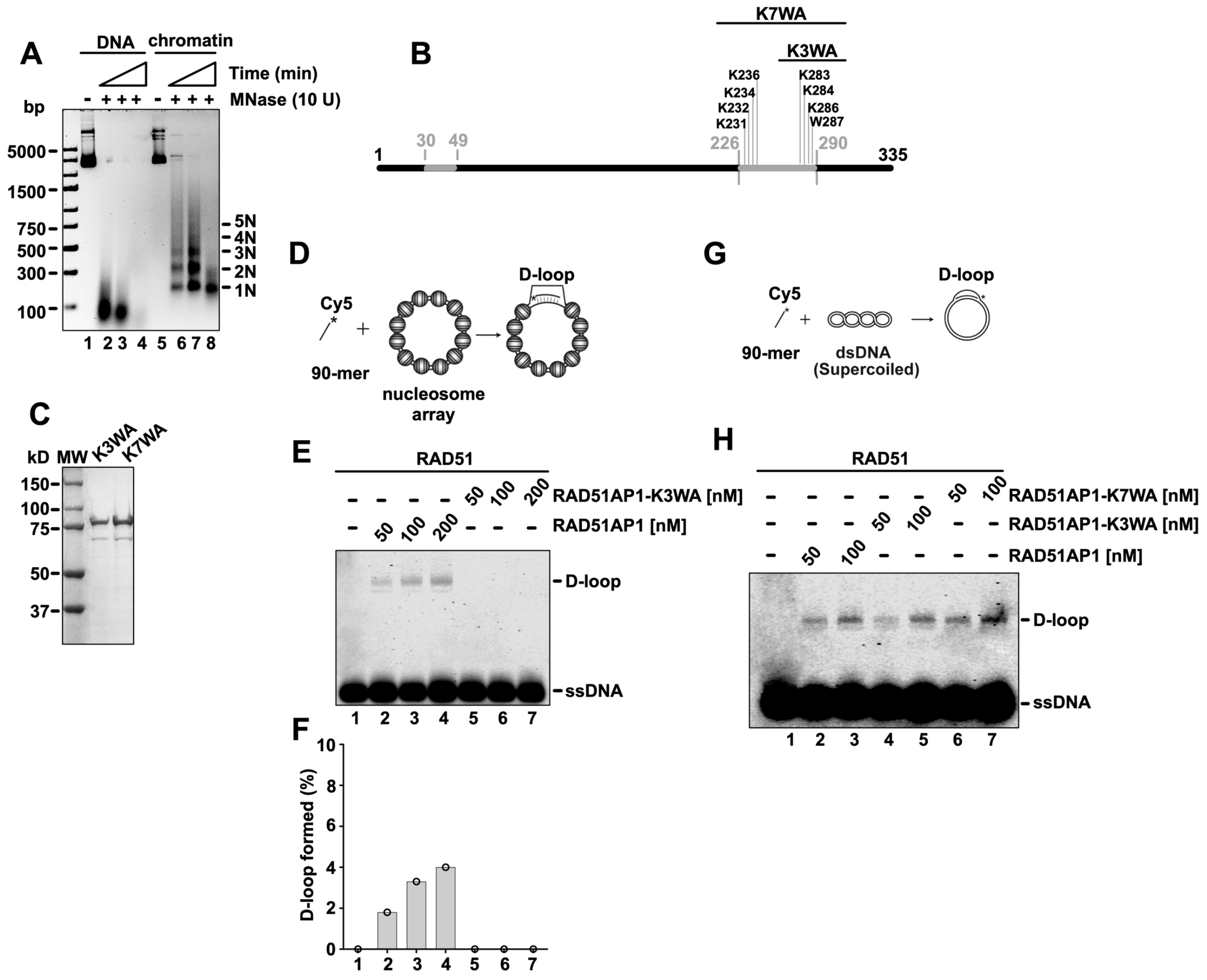


**Figure S4. RAD51AP1-K3WA and RAD51AP1-K7WA are defective in stimulating RAD51-mediated strand invasion into chromatinized DNA.** ***A***, Agarose gel obtained after limited MNase digest and deproteinization of both naked (lanes 2-4) and chromatinized pBluescript II SK(-) DNA (lanes 6-8) to show the periodicity of nucleosomes and nucleosomal repeat lengths (1N-5N). ***B***, Schematic of the full-length RAD51AP1 protein and the location of the introduced lysine/tryptophan to alanine mutations. The two DNA binding domains in RAD51AP1 are highlighted in grey. Residue numbering based on RAD51AP1 isoform 2. ***C***, SDS-PAGE of purified $His_6$-/MBP-tagged full-length RAD51AP1-K3WA and -K7WA mutants. ***D***, Schematic of the D-loop assay with chromatinized pBluescript II SK(-). ***E***, Agarose gel to show that wild type RAD51AP1 (50, 100 and 200 nM; lanes 2-4) promotes the RAD51-mediated D-loop reaction on chromatinized pBluescript II SK(-) DNA, but that RAD51AP1-K3WA (50, 100 and 200 nM; lanes 5-7) is unable to do so. ***F***, Quantification of the results in ***E*** (1 experiment). ***G***, Schematic of the D-loop assay with naked pBluescript II SK(-) DNA. ***H***, Agarose gel to show that wild type RAD51AP1, RAD51AP1-K3WA and RAD51AP1-K7WA (50 and 100 nM each) promote the RAD51-mediated D-loop reaction on naked pBluescript II SK(-) DNA.

### Supporting References

1. Dunlop, M. H., Dray, E., Zhao, W., San Filippo, J., Tsai, M. S., Leung, S. G., Schild, D., Wiese, C., and Sung, P. (2012) Mechanistic insights into RAD51-associated protein 1 (RAD51AP1) action in homologous DNA repair. *The Journal of biological chemistry* **287**, 12343-12347
